# Formulating a TMEM176B blocker in nanoparticles uncouples its paradoxical roles in innate and adaptive antitumoral immunity

**DOI:** 10.1101/2022.09.02.506404

**Authors:** Sabina Victoria, Analía Castro, Alvaro Pittini, Daniela Olivera, Sofía Russo, Ignacio Cebrian, Alvaro W. Mombru, Eduardo Osinaga, Helena Pardo, Mercedes Segovia, Marcelo Hill

## Abstract

The immunoregulatory cation channel TMEM176B plays a dual role in tumor immunity. On one hand, TMEM176B promotes antigen cross-presentation to CD8^+^ T cells by regulating phagosomal pH in dendritic cells (DCs). On the other hand, TMEM176B inhibits NLRP3 inflammasome activation through ionic mechanisms in DCs, monocytes and macrophages. Moreover, the TMEM176B blocker BayK8644 controls tumor progression through mechanisms involving inflammasome activation in prophylactic but not in therapeutic protocols. We speculated that the limited therapeutic efficacy of the compound may be linked to its potential capacity to inhibit antigen cross-presentation. Here we show that free BayK8644 inhibits antigen cross-presentation by splenic DCs. To prevent such inhibition, we reasoned that formulating BayK8644 in nanoparticles may delay the release of the compound in endosomes. Avoiding TMEM176B inhibition during the first 30 minutes of nanoparticle internalization by DCs may allow efficient cross-presentation to occur during this critical time frame. Indeed, we observed that NP-PEG-BayK8644 did not inhibit antigen cross-presentation, in contrast to the free compound. Moreover, NP-PEG-BayK8644 triggered inflammasome activation in a *Tmem176b*-dependent manner. We then injected eNP-PEG or NP-PEG-BayK8644 to mice bearing established tumors. NP-PEG-BayK8644 significantly controlled tumor growth and mice survival, as compared to eNP-PEG and free BayK8644, in a *Tmem176b*-dependent manner in mouse melanoma and lymphoma tumors. Responding animals treated with NP-PEG-BayK8644 showed reinforced tumor infiltration by total and tumor-specific CD8^+^ T cells. Overall, we rationally developed a formulating method of BayK8644 that improves its anti-tumoral therapeutic efficacy by uncoupling the dual role of TMEM176B on innate and adaptive immunity.

## Introduction

Immunotherapeutic approaches have revolutionized cancer therapy [1]. However, only a minority of treated patients with advanced disease have clinical benefits [2–4]. We thus need to identify and target novel immunoregulatory molecules to trigger effective anti-tumoral immune responses. In contrast to immune checkpoints which physiological role is to block effector immune responses, other immunoregulatory molecules such as IL-2 and interferons play dual roles, promoting and inhibiting immunity [5,6]. Such complexity is a huge challenge to therapeutically manipulate those pathways.

TMEM176B has been proposed as an emergent player in immunoregulation [7–9]. TMEM176B is an intracellular acid-sensitive non-specific cation channel which belongs to the MS4A family [10–12]. It is strongly expressed by mouse, rat and human dendritic cells (DCs) [10,12–16] among other leukocyte subsets. Functionally, TMEM176B plays a key role in tolerogenic DCs to prolong allograft survival [10]. Moreover, we have shown that TMEM176B inhibits the NLRP3 inflammasome activation through ionic mechanisms in mouse and human DCs, monocytes and macrophages [8,17,18]. Inflammasomes are cytosolic multiprotein complexes that sense cellular stress and lead to Caspase-1-dependent activation of IL-1β and IL-18 [19]. High TMEM176B expression in the tumor has been associated with diminished overall survival in patients with colon [17], glioblastoma [20] and gastric cancer [21]. Thus, TMEM176B is a potential pharmacological target in oncology. Accordingly, deletion of *Tmem176b* was associated with strong anti-tumoral immune responses in mouse cancer models that depend on host inflammasome activation and CD8^+^ T cells [17]. Moreover, pharmacological blockade of TMEM176B with BayK8644 controlled tumor growth in murine cancer models in preventive but not in therapeutic protocols [17]. We then speculated that the limited therapeutic efficacy of TMEM176B blockers may be linked to its potential capacity of inhibiting antigen cross-presentation. In fact, CD8^+^ T cells recognize tumoral antigens in the context of MHC I molecules through the cross-presentation pathway [22,23] and we have shown that *Tmem176b* promotes antigen cross-presentation by controlling phagosomal acidification in DCs [10].

In cross-presentation, endo-phagosomal antigen processing occurs through a fast kinetics, since a time limit of 25 minutes has been proposed [24]. Thus, exogenous antigens that are not processed before that time, would not be presented on MHC class I molecules [24]. We therefore speculated that formulating the TMEM176B blocker BayK8644 through a strategy that allows a slow-release kinetics in endosomes may prevent inhibition of cross-presentation by the compound but maintains its capacity to induce inflammasome activation.

Here we show that formulating BayK8644 in slow-releasing nanoparticles (NPs) triggered inflammasome activation while preventing the inhibition of antigen cross-presentation and led to improved anti-tumoral efficacy of the compound.

## Methods

### Nanoparticle Synthesis

#### I. Synthesis of CS-PEG copolymer

100 mg of chitosan-HCl (Protasan UP CL113), 27.6 mg of MeO-PEG-COOH, 2.4 mg Biotin-PEG-CO_2_H and 3.4 mg of NHS were dissolved in 14 mL of water. Then, 69 mg of EDC was added in four equal portions every 30 minutes. The resulting solution was stirred for 24 hr at room temperature, and then was ultrafiltered (Amicon 10KDa) with water and lyophilized to give chitosan-PEG as a white foam.

#### II. Preparation of CS nanoparticles

Synthesis of blank chitosan nanoparticles (NpCS) were prepared by using the solvent displacement technique, described by Torrecilla et al. [25] based on the solvent displacement method, with some modifications. Briefly, 40 mg of Lecithin were dissolved in 0.5 mL of ethanol before adding 125 μL of Crodamol and 9.5 mL of acetone. This organic phase was directly poured over 20 mL of aqueous phase containing 10 mg of CS and 50 mg of Poloxamer 188. The mixture turned milky immediately as a result of the formation of CSNp. Finally, solvents were evaporated under vacuum to a final volume of 10 mL. BayK8644 and fluorescent probe (6 coumarin) loaded CSNp formulations were prepared as described above by adding 0.5 mL of ethanolic BayK8644 solution (20 mg/mL) and 1 mL of ethanolic Coumarin solution (1mg/mL) to the organic phase respectively. The final BayK8644 concentration obtained in CSNp was 1 mg/mL.

### Encapsulation Efficiency of BayK-Loaded CSNp

The encapsulation efficiency of BayK in the CSNp was indirectly determined by the difference between the total amount of BayK in the formulation and the free drug found in the supernatant of the formulation. Accordingly, the total amount of drug was estimated by dissolving an aliquot of nonisolated BayK-loaded CSNp with acetonitrile and measured with a high performance liquid chromatography (HPLC) system. The no encapsulated drug was determined by the same method following separation of the CS structures from the aqueous medium by ultrafiltration (Amicon 10KDa). The HPLC system consisted of a Thermo Scientific Ultimate 3000 equipped with a UV detector set at 274 nm and a reverse phase Zorbax 300SB-C18 column (4.6 × 250 mm i.d., pore size 5 μm Agilent, U.S.A.). The mobile phase consisted of a 50/50 acetonitrile/buffer (0,02 % ammonium acetate solution at pH 5) mixture at a temperature of 30 ºC. The standard calibration curves of BayK were linear (r2 > 0.9998) in the range of concentrations between 0.1-1.4 μg/mL

### Animals

Six-to-ten weeks old male or female C57BL/6 Tmem176b (WT) and Tmem176b^-/-^ mice were used (Jackson Lab; Bar Harbor, ME) and bred for up to 20 generations in the Transgenic and Experimental Animals Unit (UATE) of the Pasteur Institute in Montevideo in conditions free of specific pathogens (SPF). These were kept in a controlled environment, with a temperature between 19 and 21°C, and cycles of 14 hours of light and 10 hours of darkness. Mice received water and sterile ration administered under ad libitum conditions. Tmem176b^-/-^ mice were generated in the 129/SvJ strain and heterozygous mice were backcrossed for 10 generations onto the C57BL/6 background (Janvier, Saint Berthevin, France). All experiments were performed according to local regulation and approved by the Institut Pasteur de Montevideo.

### Cell line culture

The E.G7-OVA (expressing OVA antigen) cell line (ATCC® CRL-2113™, (Manassas, VA), thymic lymphoma of murine origin) was cultured in DMEM medium supplemented with 10% fetal bovine serum (SBF), 10 mM HEPES, 1 mM sodium pyruvate, MEM-1% non-essential amino acids, 0.05 mM β-mercaptoethanol, penicillin/streptomycin 100 units/mL and 0.4 mg/mL geneticin. It was kept at 37°C in the presence of 5% CO_2_.

### Primary culture of dendritic cells derived from bone marrow (BMDCs)

Dendritic cells derived from bone marrow (BMDCs) were obtained. For this, the femurs and tibiae bone marrow were extracted of Tmem176b and *Tmem176b*^*-/-*^ C57BL/6 mice. After red blood cells lysis, 5.0×10^6^ cells were seeded in Petri dishes, in RPMI supplemented medium (10% fetal bovine serum, 10 mM HEPES, 1 mM sodium pyruvate, 1 % non-essential MEM-amino acids, 2 mM L-glutamine, 0.05 mM β-mercaptoethanol, penicillin/streptomycin 100 units/mL). Thus, bone marrow-derived DCs were differentiated by culturing bone marrow cells for 8 days in the presence of 0.4 ng/mL GM-CSF. On day 3 and 6, the growth medium was renewed, adding GM-CSF to the culture each time. At day 8, adherent cells were >95% CD11c^+^CD11b^+^MHC II^int^ evaluated by flow cytometry analysis.

### Nanoparticles uptake by Bone Marrow Dentritic Cells (BMDCs)

At day 8, BMDCs were detached with a buffer solution containing SFB and EDTA. For confocal microscopy internalization studies, 1.0×10^5^ cells were seeded in a 12-well plate, glass cover slips were previously placed in each well. Cells were allowed to adhere for 1,5 hours and stimulated for 30 minutes with NP-PEG-6cou at 37°C to allow endocytosis process or 0°C (as a negative control), washed and then chased for 15 minutes at the same conditions. All images were taken using a Zeiss LSM 800™ semi-spectral confocal laser scanning microscope (Beckman Coulter). For flow cytometry internalization studies, the same protocol was followed as mentioned above. BMDCS were collected and samples were acquired using Cyan ADP Flow Cytometer equipped with a 488 nm laser and a band filter (530/40 nm). For data analysis FlowJo v.X software (BD, Ashland, OR, USA) was used.

### Colocalization Studies by Confocal Microscopy

*Tmem176b*^*+/+*^ (WT) and *Tmem176b*^*-/-*^ C57BL/6 BMDCs were plated in 12-well plate (1×10^5^ cells/well) containing glass cover slips and cultured for 1,5 hours. Then cell were treated with NP-PEG-6-cou during 30 minutes, washed and chased for 15 minutes. Cells were fixed in 2% (w/v) paraformaldehyde (PFA) for 15 min, permeabilized and blocked using TritonX100 0,1%, 3% BSA in PBS. Primary antibody, rabbit anti-mTMEM176B Polyclonal antibody (Proteintech Group) 5 μg/mL, or a control IgG antibody, was diluted in BSA-PBS-Triton solution and incubated overnight at 4 °C. Three washing steps were performed using PBS solution. Biotina-donkey anti-Rabbit IgG (H+L) secondary antibody diluted 1:500 in BSA-PBS-Triton was added and incubated for 1,5 hours. After washing, cells were incubated with Streptavidin Alexa Fluor 568 conjugate (Molecular Probes) diluted 1:1000 in BSA-PBS-Triton. Finally, nuclei were stained with DAPI diluted 1:5000 (Life Technologies) and incubated for 5 minutes. Sections were mounted in ProLong antifade mounting medium (Molecular Probes). Fluorescence was observed by confocal microscopy Zeiss LSM 800™ semi-spectral.

### *In Vitro* Inflammasome Activation

IL-1β was quantified in BMDCs culture supernatant as a readout of the NLRP3 inflammasome activation. As mentioned above, at day 8, 5.0×10^4^ BMDCs were seeded in a 96-well plate, and allowed to adhere for 1,5 hours. Then, cells were stimulated for 3 hs with 0.25 μg/mL LPS, washed and treated for 2 hours with the indicated doses of free BayK8644, NP-PEG-BayK8644 and eNP-PEG. The presence of IL-1β was determined with a ELISA kit, according to the manufacturer protocol (Biolegend, 432603).

To determine Caspase-1 activation by Western blot, culture supernatants from BMDCs stimulated in the absence of FBS were precipitated with 20%(v/v) TCA and washed with acetone. Cell lysates were generated with RIPA buffer in the presence of a protease inhibitor cocktail. Cell lysates and precipitates from culture supernatants were electrophoresed, blotted and probed with anti-Caspase-1 (Adipogen, AG-20B-0042) antibodies. Quantification was carried out using ImageJv.1.5i software and for statistical analysis.

Caspase-1 activation was quantified in Tmem176b BMDCs culture as a readout of the NLRP3 inflammasome activation. As mentioned above, at day 8, BMDCs were seeded (5×10^5^cells) in 35 mm culture dishes, and allowed to adhere for 1,5 hours. Cells were stimulated for 2 hr with 20 μM of free BayK8644, NP-PEG-BayK8644 and eNP-PEG and then washed. BMDCs were stained with FLICA-1 (FLICA 660 Caspase-1 Assay, ImmunoChemistry) for 30 minutes, and analyzed by flow cytometry using Cyan ADP Flow Cytometer equipped with a 488 nm laser and a band filter (530/40 nm). For data analysis FlowJo v.X software (BD, Ashland, OR, USA) was used.

### Antigen cross-presentation assays in mouse splenic dendritic cells

To obtain splenic DCs, spleen of C57Bl6 mice was extracted, cut into small pieces and treated with collagenase-D (Roche, Merck). After 30 minutes of incubation at 37 ° C, the enzymatic reaction was stopped with a 10 mM EDTA solution. Lysis of red blood cells was performed with an ammonium chloride solution. Finally, cells were washed with PBS/SFB1%. For the isolation of CD11c^+^ cells, murine anti-CD11c monoclonal antibodies coupled to magnetic microspheres supplied in the commercial kit “CD11c MicroBeads UltraPure, mouse” (Miltenyi Biotec Inc, Auburn, CA, USA) were used according to the protocol suggested by the manufacturer. Briefly, splenic cells were labeled with anti-CD11c antibodies conjugated to the beads. The cells were allowed to pass through a magnetic column subjected to a magnetic field. Thereby, CD11c^+^ cells were retained on the column. Then, the column was removed from the influence of the magnetic field and cells were eluted. The purity of the CD11c^+^ fraction was confirmed by flow cytometry.

CD11c^+^ spleen cells were incubated with 3 mg/mL of soluble OVA, or OVA plus 20 μM of free BayK8644, 20 μM of BayK8644 encapsulated into chitosan nanoparticles (NP-PEG-BayK8644) or empty counterpart (eNP-PEG) for 4 hours in supplemented-RPMI medium. Also, as a control, CD11c^+^ spleen cells were incubated with SIINFEKL peptide at indicated concentrations or SIINFEKL peptide plus BayK8644 for 4 hours in supplemented-RPMI medium. Cells were then washed with PBS/BSA 1% and fixed with 0.008 % glutaraldehyde for 3 minutes on ice and quenched with 0.2 M glycine solution. After one final wash with PBS, CD11c^+^ spleen cells were incubated with the B3Z cell line (T cell hybridoma cell line) for 18 hours at 37°C and 5% CO_2_. B3Z cell line specific for SIINFEKL allowed us to study the T cells activation, and was kindly provided by Ignacio Cebrián (Membrane Fusion Laboratory, IHEM-CONICET; Faculty of Medical Sciences, UNCuyo). The B3Z cell line is a β-galactosidase (lacZ) inducible T cell hybridoma. They are stably transfected with the Dectin-1-CD3ζ chimeric construct (Dectin-1 extracellular region fused to the CD3ζ cytoplasmic region). Furthermore, these cells are transiently transfected with an NFAT-LacZ reporter gene (activated T-cell nuclear factor NFAT, fused to the E. coli lacZ gene); since CD3ζ triggers an intracellular signal that leads to the activation of the transcriptional factor NFAT. Previous studies demonstrated that the heterologous lacZ gene, under the control of the complete enhancer region of IL-2 or NFAT alone, is specifically induced in transfected and activated T cells. Therefore, activation of transfected T cells results in the synthesis of the products of the IL-2 and lacZ genes. The activation of B3Z cell line was measured detecting β-galactosidase activity by optical density (absorbance at 595–655 nm) using chlorophenol red-β-D-galactopyranoside (CPRG; Roche) as substrate for the reaction.

### *Ex vivo* isolation and flow cytometry analysis of CD8+ T cells

Tmem176b C57BL/6 mice were injected s.c with 0,5 × 10^6^ EG7 thymic lymphoma cells. When tumors were palpable (tumor size of 5mm x 5mm), 70 nmoles of BayK8644 encapsulated in NPs was intratumorally injected. NPs injections were repeated at the +2, +4 days after the first injection. After the third NP-PEG-BayK8644 injection, tumors were resected.

Animals treated with NP-PEG-BayK8644 were classified according to tumor growth kinetics. Those animals that control tumor growth over time (tumor size at the resection time were <100mm^2^) were named Responders. Inversely, those animals did not control the tumor size were called Progressors. T cells were isolated by dissociating tumor tissue in the presence of collagenase D (1 mg/ml) (Roche-Merck) for 10 min at 37°C. Reaction was stopped with EDTA 10mM, and the suspension was filter through a 50 μm strainer. After red blood cell lysis, flow cytometry analysis was performed. For flow Cytometry analysis, single cell suspensions were stained with antibodies against surface molecules CD8α-APC-Cy7, TCRβ-PE, CD19-PercPCy5.5, OVA-pentamer-APC and TCRvβ12-FITC to discriminate EG.7-OVA tumor cells. All data were collected on a FACS Aria Fusion (BD Biosciences) and analyzed with FlowJo software (FlowJo v.X software (BD, Ashland, OR, USA).

### Tumor Models and Treatments

*Tmem176b*^*+/+*^ (WT) and *Tmem176b*^*-/-*^ C57BL/6 mice were injected s.c with 0.5 × 10^6^ E.G7-OVA thymic lymphoma cells or 0.5 × 10^5^ 5555 melanoma cells. At the indicated experiment, i.t injection was performed alternating one WT and one *Tmem176b*^*-/-*^ mouse until completing both groups. In treated animals, alternation was done between BayK8644- and vehicle-treated animals or NP-PEG-BayK8644 and eNP-PEG-treated animals. Tumor growth was caliper measured manually every 2-3 days, and the two major diameters were taken. 70 nmoles of free BayK8644 or BayK8644 encapsulated in chitosan nanoparticles, or the same amount of empty NPs injections were repeated at the +2, +4, +6, +8 days after the first injection. Mice were sacrificed when one of the major tumor diameters reached 2 cm.

### Statistical analysis

Statistical analyses were performed by GraphPad Prism 7 (GraphPad Software, Inc., La Jolla, CA). Survival analyses were done with the Log-rank (Mantel-Cox) test. Comparison of two experimental conditions was done with unpaired Student’s t test. Comparison of multiple conditions was done with one-way ANOVA tests. Differences in gene expression between responder and progressors groups were assessed using the unpaired t-test when normality assumption was met.

## Results

### Encapsulation of BayK8644 in chitosan nanoparticles

We first studied whether BayK8644 inhibits antigen cross-presentation *in vitro*. Enriched splenic CD11c^+^ DCs from WT mice were pulsed with OVA protein and co-cultured with the OVA-specific CD8^+^ B3Z T cell hybridoma in which the β-Gal enzyme is under the control of the IL-2 promoter [26]. OVA induced a dose-dependent activation of B3Z cells **(Supplementary figure 1)**. We observed that treating splenic DCs with BayK8644 significantly decreased their capacity to activate B3Z cells **(Figure 1A)**. Conversely, BayK8644 did not inhibit activation of B3Z cells induced by the minimal OVA peptide SIINFEKL **(Figure 1B)**. Thus, these results suggest that Bayk8644 interferes with the processing of OVA protein through the cross-presentation pathway in splenic DCs as expected for a TMEM176B inhibitor [10]. This observation may explain the limited therapeutic efficacy of the compound [17].

**Figure 1.**
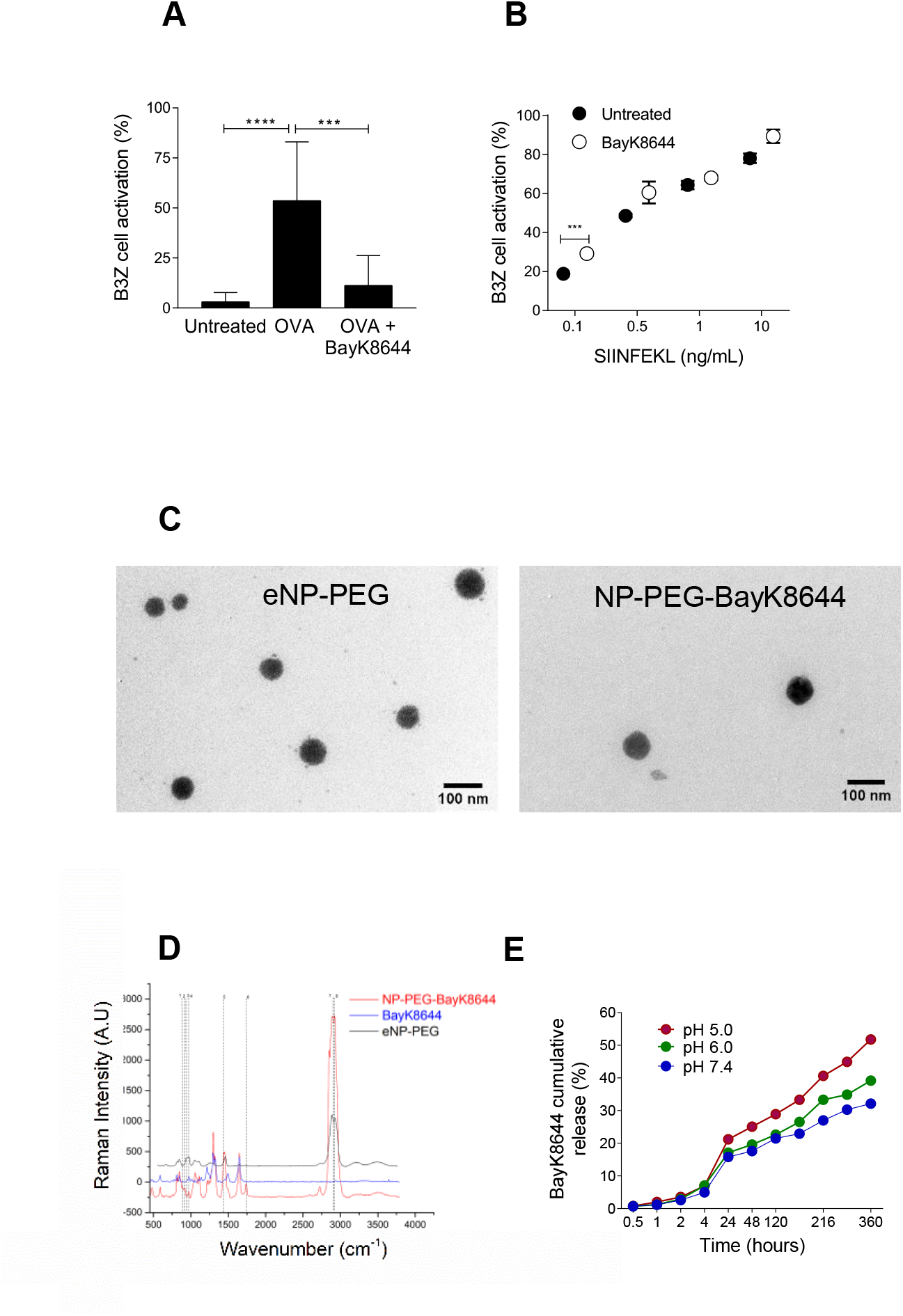
Encapsulation of BayK8644 in chitosan nanoparticles. **a)** CD11c^+^ splenic DCs were loaded with 3mg/mL soluble OVA for 4 hr ± 20 μM of free BayK8644. Then, DCs were extensively washed, fixed, and cultured (18 hs) with the β-galactosidase-inducible B3Z T-cell hybrid, which is specific for the OVA peptide (SIINFEKL) in association with H-2K^b^ MHC-I molecules. B3Z T cell activation was measured by assessing β-galactosidase activity in a colorimetric assay. A pool of three independent experiments is shown. ***p < 0.001, ****p < 0.0001. One-way ANOVA test. **b)** Splenic DCs were loaded for 45 minutes with the OVA peptide SIINFEKL, which does not need further processing in intracellular compartments to be presented on H[2K^b^ MHC[I molecules ± 20μM BayK8644. DCs were then washed and fixed and co-cultured with B3Z T cells. A pool of three independent experiments is shown. ***p < 0.001. Unpaired Student’s *t* test. **c)** Transmission electron micrograph (TEM) image showing spherical morphology and physical diameter of NPs. Note that NPs sizes are fairly uniform and it reveals the conserved NPs size distribution. **d)** The Raman spectrum of NP-PEG-BayK (in red), free BayK8644 (blue) and eNP-PEG (black). **e)** Cumulative drug release profiles of NP-PEG-BayK8644 at different pH conditions. The NP-PEG-BayK8644 displayed a pH-dependent BayK8644 release profile. NP-PEG-BayK8644 solution was dissolved in RPMI culture media at the indicated pH (Ph 5.0, pH 6.0 and pH 7.4). The amount of free BayK8644 released was determined by HPLC at different time points.

We then reasoned that formulating BayK8644 through a slow releasing mechanism in the endosomal compartment may avoid inhibition of cross-presentation by the compound. Chitosan nanoparticles (NPs) have been characterized as drug-encapsulating strategies that release compounds with a slow kinetics [27,28]. We therefore speculated that encapsulating BayK8644 in chitosan NPs may avoid inhibition of cross-presentation by the compound while probably keeping its inflammasome-activating capacity. Chitosan NPs (semi-crystalline linear polysaccharide composed of N-acetyl D-glucosamine and D-glucosamine units linked by β (1-4) glucosidic bonds) were synthesized and coated on their surface with polyethylene glycol (PEG), which forms a protective hydrophilic layer around the nanoparticles [25]. Thus, empty pegylated NPs (eNP-PEG), pegylated NPs containing the compound (+)-BayK8644 (NP-PEG-BayK8644) and pegylated NPs containing the fluorescent probe 6-coumarin (NP-PEG-6cou) were synthesized. The average size, the polydispersity index (PI) and the Z potential of the NPs, as well as the efficiency of encapsulation of the formulation containing BayK8644, are detailed in **Supplementary table 1**. The NPs size was obtained by using a Nano Zetasizer, showing an average size close to 200 nm, which is within the range of optimal sizes to improve the biological fluids circulation time of NPs, as well as for its cellular internalization [29]. We can also conclude that the colloidal solutions obtained are stable, since their Z potential is less than 30 mV. The BayK8644 encapsulation efficiency in all the performed syntheses was always greater than 99%. The TEM images of NP-PEG-BayK8644 and eNP-PEG show their spherical shape and a homogeneous size distribution **(Figure 1C)**. The sizes obtained by TEM are smaller than those obtained by Nano Zetasizer and shown in **Supplementary table 1**. This difference in size measurement arises because in solution the ions of opposite charge will be attracted to the surface of the nanoparticles generating ionic layers that move along with it, and the Nano Zetasizer measures the nanoparticles hydrodynamic diameter in solution, while the TEM reports the actual diameter of the NPs without the ionic layer [30].

The study by confocal Raman microscopy allowed us to show that there are notorious differences between the spectra of pure BayK8644 and chitosan NPs with respect to that obtained with chitosan NPs loaded with BayK8644, obtaining a spectrum different from the one that would have been obtained only by adding the individual spectra **(Figure 1D)**. This indicates that there must have been a chemical interaction between the constituent materials of the NPs and the BayK8644 molecule. The latter is important when it comes to obtaining an adequate release of the BayK8644 molecule.

We then studied the *in vitro* release kinetics of BayK8644 from NPs at different pHs. As expected [27,28], we observed a slow kinetics release of the compound **(Figure 1E)**. Acidic pH increased the percentage of released BayK8644 since 24 hr incubation of the NPs with the buffer **(Figure 1E)**. Thus, encapsulation of BayK8644 in chitosan NPs is highly efficient, generates NPs with physicochemical characteristics compatible with their use with cells and releases the compound with a slow kinetics.

### NP-PEG-BayK8644 do not inhibit antigen cross-presentation by DCs but still trigger inflammasome activation

To further determine whether encapsulation of BayK8644 in chitosan NPs may reinforce the anti-tumoral properties of the compound, we studied whether and how NP-PEG-BayK8644 modulate DCs biology. First, we assessed their internalization by DCs. Flow cytometry studies showed that bone marrow-derived DCs (BMDCs) efficiently internalized NP-PEG-6cou, increasing both the percentage of 6-coumarin^+^ cells **(Figure 2A, B)** and 6-coumarin mean fluorescence intensity (MFI) **(Figure 2A, C)** in comparison to untreated cells and BMDCs incubated at 0°C as a control to minimize active internalization. Confocal microscopy experiments showed that at 0°C the NPs were mostly associated to the cell surface, whereas at 37 °C the NPs were fully internalized by BMDCs **(Figure 2D)**. Moreover, we observed that NP-PEG-6-cou were localized within Tmem176b^+^ compartments **(Figure 2E)**, likely endosomes [10]. Thus, BayK8644 released by NPs within endosomes may be in proximity to Tmem176b.

**Figure 2.**
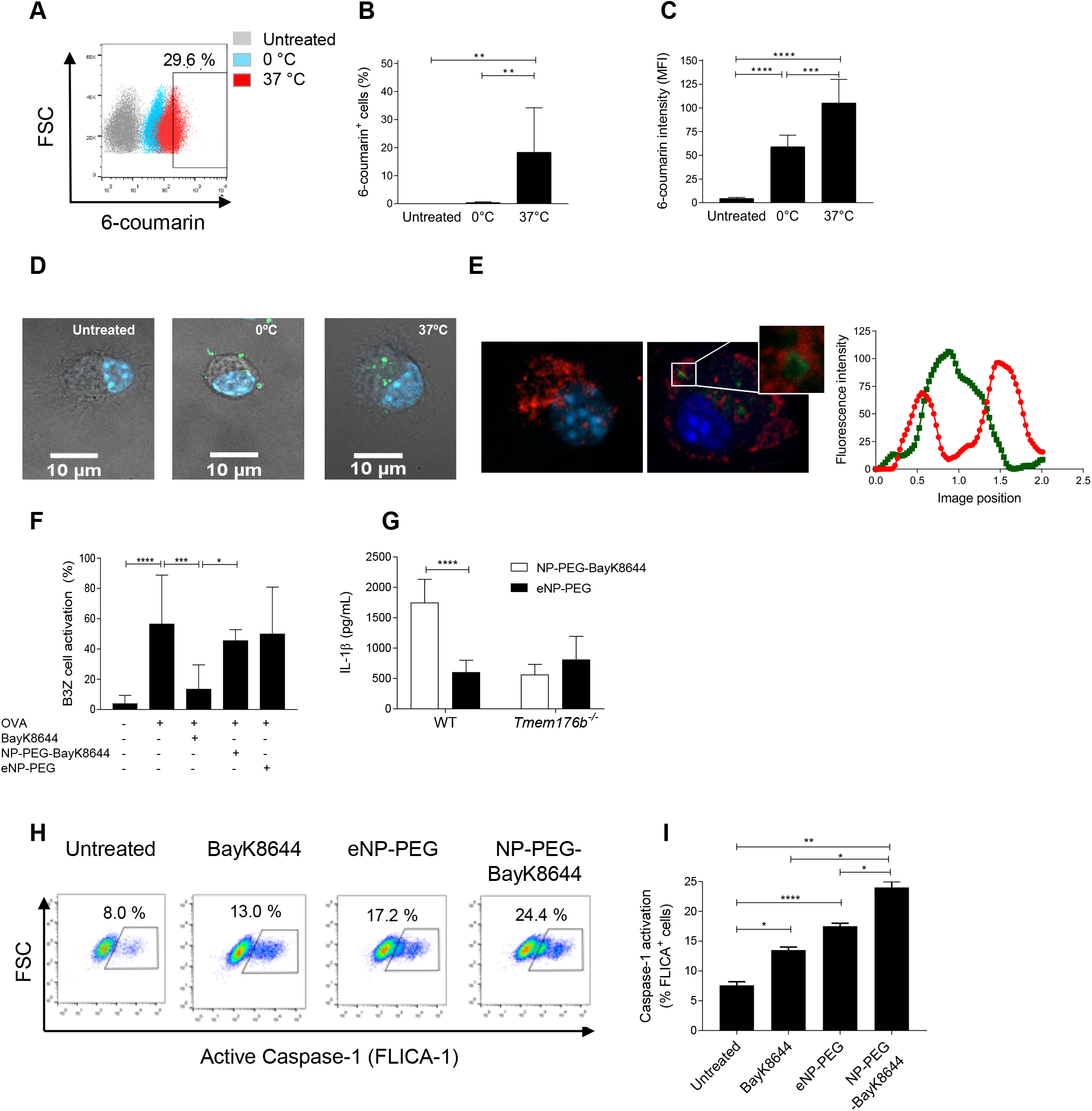
Chitosan nanoparticles are internalized by DCs, do not inhibit antigen cross-presentation and trigger inflammasome activation. **a)** BMDCs were treated *in vitro* with 6-coumarin encapsulated NPs (NP-PEG-6cou) for 30 minutes, at 37 °C or 0 °C as a negative endocytosis control. A representative dot plot is shown with the percentage of 6-coumarin positive cells of BMDCs treated with NP-PEG-6cou at 37°C (red dots), compared with BMDCs treated with 6-coumarinNP-PEG at 0°C (cyan dots). Grey dots represent untreated BMDCs. One experiment representative of three is shown. **b)** Percentage of 6-coumarin positive cells. A pool of three independent experiments is shown. ** p < 0.01, One-way ANOVA test. **c)** Mean Fluorescence Intensity (MFI) of 6-coumarin dye determined by flow cytometry. A pool of three independent experiments is shown. *** p < 0.001; ****p < 0.0001. One-way ANOVA test. **d)** Confocal microscopy analysis of BMDCs treated with NP-PEG-6cou (green fluorescence) for 30 minutes at the indicated temperatures (0°C and 37°C). Nuclei were stained with DAPI (blue fluorescence). One experiment representative of three is shown. At least 50 cells were analyzed/experiment. **e)** Confocal microscopy study showing the detection of NP-PEG-6cou and the endogenous Tmem176b protein. BMDCs from WT mice were treated with NP-PEG-6cou for 30 minutes, then immunostaining was performed with anti-Tmem176b (red) antibody. Nuclei were stained with DAPI (blue). The graph shows the intensity profiles of anti-Tmem176b staining (red) and NP-PEG-6cou (green) obtained using ImageJ software, along the indicated yellow line inside the white square. The fluorescence intensities are plotted along the y-axis and the image position along the x-axis. One experiment representative of three is shown. At least 50 cells were analyzed/experiment. **f)** CD11c^+^ splenic DCs were left untreated or loaded with 3mg/mL soluble OVA for 4 hr ± 20 μM BayK8644, eNP-PEG or NP-PEG-BayK8644. B3Z T cell activation was assessed through a colorimetric assay. A pool of three independent experiments is shown. *p<0.05, ***p < 0.001, ****p < 0.0001. One-way ANOVA test. **g)** WT and *Tmem176b*^*-/-*^ BMDCs were treated with LPS (0.25 μg/mL) for 3 hr, washed and exposed for 2 hr to 50 μM BayK8644 encapsulated in chitosan nanoparticles (NP-PEG-BayK8644), or an equivalent amount of empty NPs (eNP-PEG). IL-1β was measured in the culture supernatant by ELISA. One experiment representative of three is shown. ****p < 0.0001. Two-way ANOVA test. **h)** WT BMDCs were treated with LPS (0.25 μg/mL) for 3 hr, washed and exposed for 2 hr to 50 μM BayK8644 encapsulated in chitosan nanoparticles (NP-PEG-BayK8644), or an equivalent amount of empty NPs (eNP-PEG) or 50 μM free BayK8644. Inflammasome activation was studied by assessing Caspase-1 activation by flow cytometry using the FLICA-1 reagent. Representative dot plots are shown for the FLICA quantification by flow cytometry. One experiment representative of five is shown. **i)** A pool of the five experiments commented in H is shown. *p<0.05, **p<0.01, ****p<0.0001. One-way ANOVA test.

We then wished to determine whether BayK8644 encapsulated in NPs may inhibit antigen cross-presentation by DCs. Splenic DCs were loaded with OVA protein ± BayK8644 or ± NP-PEG-BayK8644 and co-cultured with B3Z CD8^+^ T cells. We observed that in contrast to free BayK8644, NP-PEG-BayK8644 failed to block activation of B3Z cells **(Figure 2F)**. Furthermore, treatment of DCs with NP-PEG-BayK8644 induced the secretion of IL-1β at higher levels than eNP-PEG in WT but not in *Tmem176b*^*-/-*^ DCs **(Figure 2G)**. Moreover, NP-PEG-BayK8664 were more effective than eNP-PEG and BayK8644 in inducing Caspase-1 activation in DCs **(Figures 2H-I and Supplementary figure 2)**. Thus, NP-PEG-BayK8644 trigger inflammasome activation in DCs while preventing inhibition of antigen cross-presentation by BayK8644.

### Therapeutic injection of BayK8644 encapsulated in chitosan NPs increases survival and retards tumor growth in a *Tmem176b*-dependent manner

Having shown that NP-PEG-BayK8644 do not inhibit antigen cross-presentation **(Figure 2F)** while still triggering inflammasome activation **(Figures 2G-J and Supplementary figure 2)**, we then studied whether those NPs may control tumor growth *in vivo* in a therapeutic setting. We had shown that although BayK8644 injected i.p before tumor establishment controlled its growth (preventive protocol), it failed to do so in animals with established tumors (therapeutic protocol) [17].

To evaluate the therapeutic efficacy of NP-PEG-BayK8644, *in vivo* studies were carried out in WT mice carrying established 5555 and EG7 tumors. We observed that intratumoral injection of NP-PEG-BayK8644 significantly improved mouse survival in melanoma **(Figure 3A)** and lymphoma **(Figure 3B)** models as compared to animals injected with eNPs-PEG. In contrast, when *Tmem176b*^*-/-*^ animals were treated with NP-PEG-BayK8644 or eNP-PEG, we found no difference in mouse survival **(Figure 3C)**. Thus, NP-PEG-BayK8644 control tumor growth through a *Tmem176b*-dependent mechanism. Additionally, intratumoral injection of free BayK8644 also failed to control tumor growth and to improve mouse survival in comparison to vehicle-treated animals **(Figure 3D)**. To further understand how NP-PEG-BayK8644 may control tumor growth, we assessed inflammasome activation in the tumor-draining lymph nodes (TDLN) of animals treated with NP-PEG-BayK8644 or eNP-PEG. In fact, intratumorally injected fluorescent NPs were found in the TDLN 24h post-administration in resident but not in migratory conventional (co)DCs **(Supplementary figure 3)**. We did not observe different percentages of active Caspase-1 (FLICA-1+) cells within resident coDCs in animals injected with eNP-PEG versus NP-PEG-BayK8644 **(Figure 3E)**. Moreover, in mice injected with NP-PEG-BayK8644, progressor and responder animals showed no differences in Caspase-1 activation within resident coDCs **(Figure 3F)**. Thus, control of tumor growth in mice injected with NP-PEG-BayK8644 versus eNP-PEG may not be explained by differential capacity to trigger inflammasome activation.

**Figure 3.**
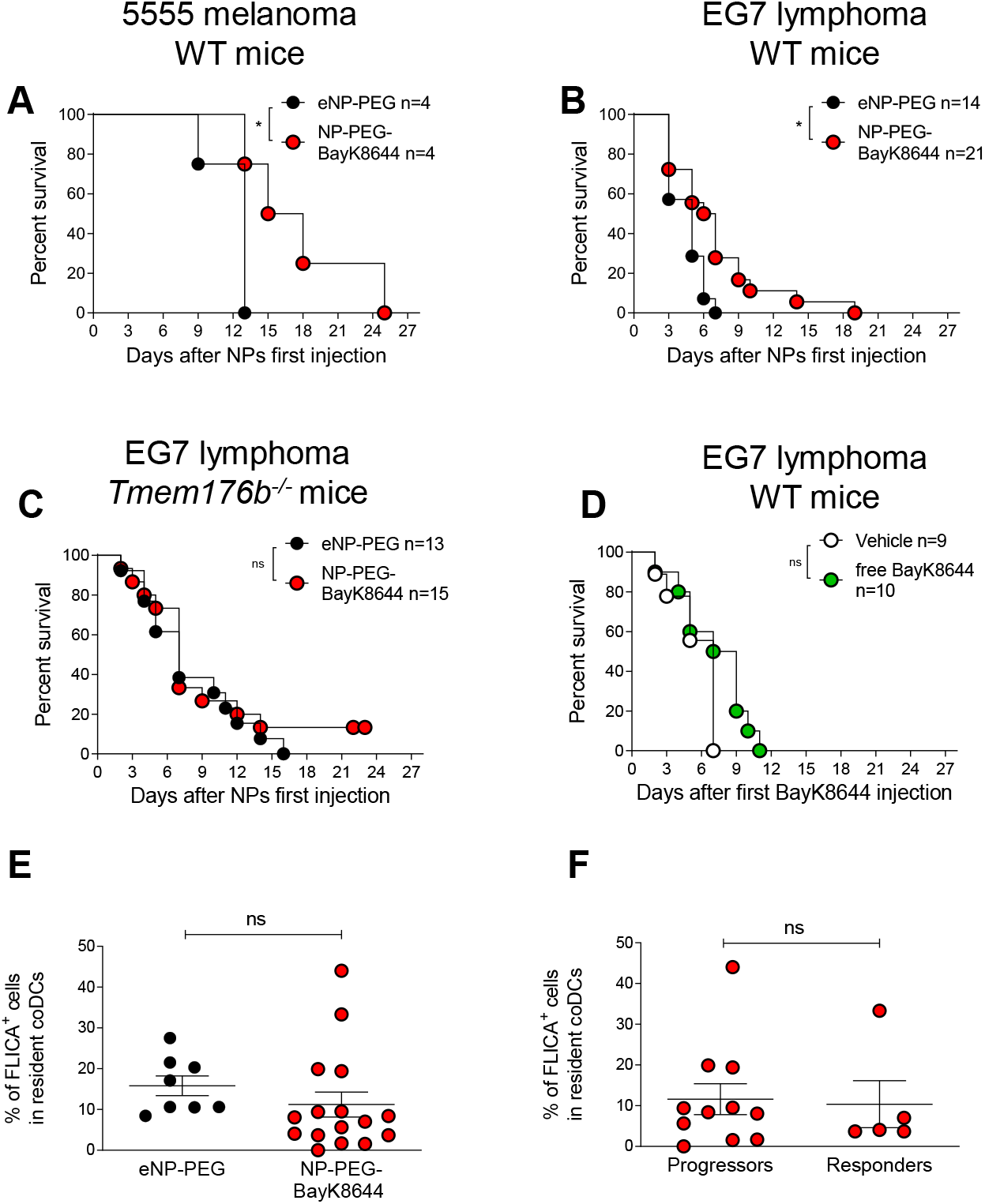
Therapeutic treatment with BayK8644 encapsulated in chitosan nanoparticles (NP-PEG-BayK8644) increases mice survival and reduces tumor growth compared with empty nanoparticles (eNP-PEG). WT and *Tmem176b*^*-/-*^ animals were inoculated with 5555 melanoma or EG7 lymphoma tumors. Once tumors reached 25 mm^2^, animals were i.t. injected with 70 ×10^−9^ moles of BayK8644, NP-PEG-BayK8644 or eNP-PEG. Treatments were repeated every two days reaching a total of five injections. Mice were euthanized when the longer diameter reached tumors reached >200 mm. **a)** Mouse survival of WT mice bearing 5555 melanoma and injected with NP-PEG-BayK8644 or eNP-PEG. *p< 0.05. Log-rank (Mantel-Cox) Test. **b)** Mouse survival of WT mice bearing EG7 lymphoma and injected with NP-PEG-BayK8644 or eNP-PEG. *p< 0.05. Log-rank (Mantel-Cox) Test. **c)** Mouse survival of *Tmem176b*^*-/-*^ mice bearing EG7 lymphomas and injected with NP-PEG-BayK8644 or eNP-PEG. ns= not significant. Log-rank (Mantel-Cox) Test. **d)** Mouse survival of WT mice injected with vehicle control or free BayK8644. ns= not significant. Log-rank (Mantel-Cox) Test. **e)** Percentage of FLICA-1 (Active Caspase-1) positive cells within resident coDCs in the TDLN 3 days after the third NP injection in EG tumor-bearing WT animals. ns= not significant. Student’s *t* test. **f)** Percentage of FLICA-1 (Active Caspase-1) positive cells within resident coDCs in the TDLN 3 days after the third NP-PEG-BayK8644 injection in EG tumor-bearing WT animals. Comparison of progressors and responder animals. ns= not significant. Student’s *t* test.

### Increased tumor infiltration by total and tumor-specific CD8^+^ T cells is associated with tumor control in NP-PEG-BayK8644-treated animals

We then speculated that preventing antigen cross-presentation inhibition by BayK8644 may enhance the anti-tumoral efficacy of NP-PEG-BayK8644. Anti-tumoral CD8^+^ T cells are primed in the TDLN through the cross-presentation pathway [23]. We therefore studied CD8^+^ T cells by flow cytometry in the TDLN form EG7 tumors-bearing animals treated with eNP-PEG, NP-PEG-BayK8644 or free BayK8644. According to our hypothesis, absolute numbers of total and tumor-specific CD8^+^ T cells were increased in the TDLN from NP-PEG-BayK8644 mice in comparison with BayK8644 and eNP-PEG-treated animals, although statistical significance was not reached **(Figure 4A-B)**. A similar trend was found in the tumor microenvironment for absolute numbers of total and tumor-specific **(Figure 4C-D)** CD8^+^ T cells, when comparing the three therapeutic groups. Then, we found a significant negative correlation between the size of tumors and the absolute numbers of total **(Figure 4E)** and tumor-specific **(Figure 4F)** CD8^+^ T cells within the NP-PEG-BayK8644-treated group in the tumor microenvironment. Accordingly, we observed a significant increase in the percentage and absolute number of intratumoral total CD8^+^ T cells in animals controlling tumor growth (responders) versus progressors within the NP-PEG-BayK8644-treated group **(Figure 4G-I)**. Moreover, similar results were found when analyzing intratumoral tumor-specific CD8^+^ T cells **(Figure 4J-L)**. Our results therefore show an association between clinical responses and tumor infiltration by CD8^+^ T cells, supporting a role for preserved antigen cross-presentation in the anti-tumoral properties of NP-PEG-BayK8644.

**Figure 4.**
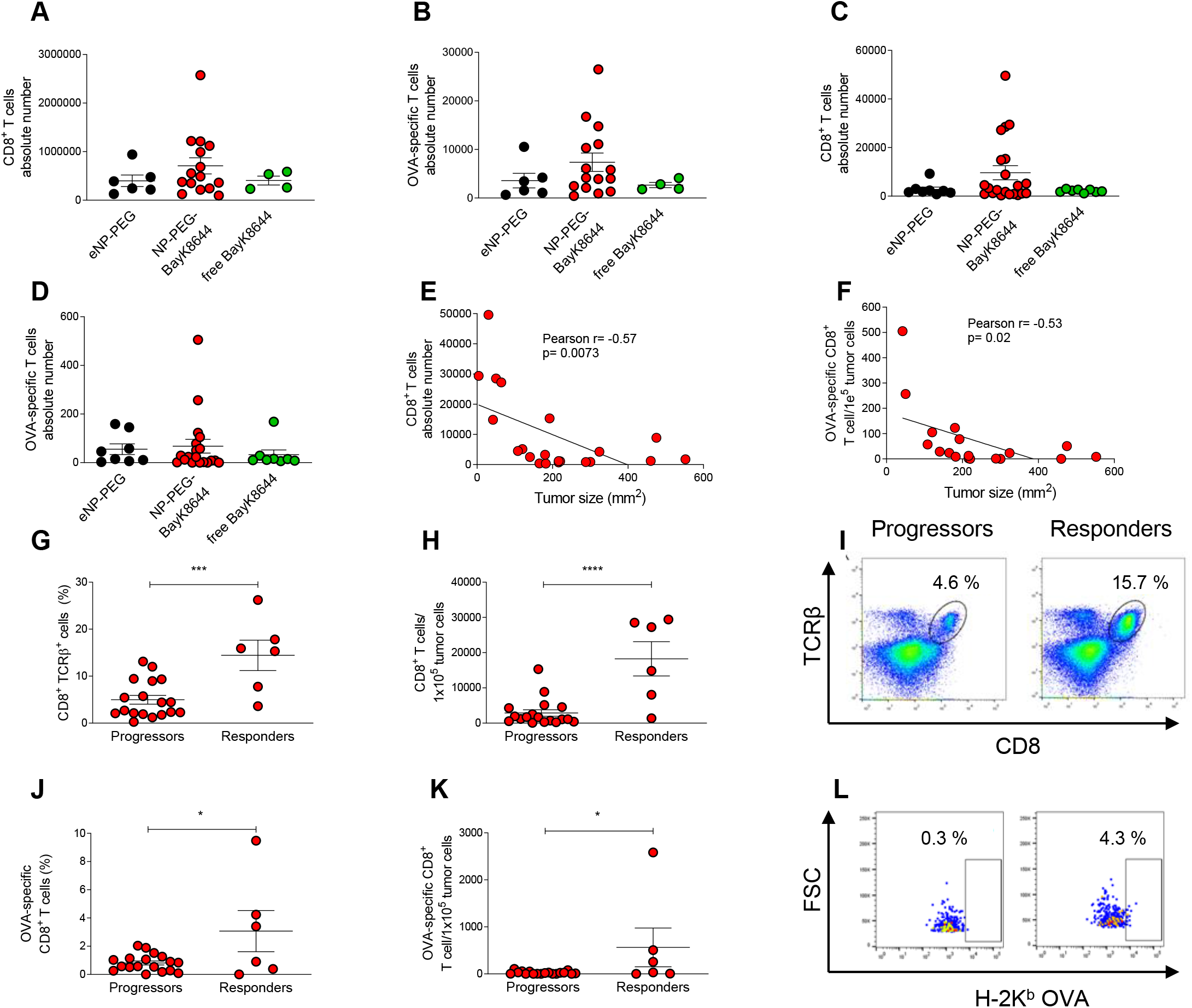
Tumor control in NP-PEG-BayK8644-treated mice is associated with tumor cell infiltration by CD8^+^ T cells. WT mice were subcutaneously injected with 5.0×10^5^ cell line E.G7-OVA. BayK8644, eNP-PEG or NP-PEG-BayK8644 were i.t injected every 2 days when tumors size reached 25 mm^2^. CD8^+^ T cells were studied in cellular suspensions from TDLNs and tumors 24 hs after the third NP or BayK8644 injection from the indicated groups. Animals from three experiments are shown. Absolute numbers are expressed by 1×10^5^ tumor cells (Vβ12^+^ cells). When indicated, NP-PEG-BayK8644-treated animals were classified as responders if at the resection time their tumor sized <100mm^2^ and as progressors if they sized >100 mm^2^. **a)** Absolute numbers of total CD8^+^ T cells in tumor-draining lymph nodes (TDLN). **b)** Absolute numbers of OVA-specific CD8^+^ T cells in TDLN. **c)** Absolute numbers of total CD8^+^ T cells in tumors. **d)** Absolute numbers of OVA-specific CD8^+^ T cells in tumors. **e)** Correlation study between the tumor-infiltrating total CD8^+^ T cell absolute numbers and tumor size (mm^2^). Correlation test. **f)** Correlation study between the tumor-infiltrating OVA-specific CD8^+^ T cell absolute numbers and tumor size (mm^2^). Correlation test. **g)** Relative frequency of total CD8^+^ T cells among Vβ12^-^ cells in tumors of NP-PEG-BayK8644-treated mice comparing progressor and responder animals. *** p < 0.001, Student’s *t* test. **h)** Absolute numbers of total CD8^+^ T cells among Vβ12^-^ cells in tumors of NP-PEG-BayK8644-treated mice comparing progressor and responder animals. **** p < 0.0001, Student’s *t* test. **i)** Representative flow cytometry dot plots of total CD8^+^ T cells in tumors of NP-PEG-BayK8644-treated mice comparing progressor and responder animals. **j)** Relative frequency of OVA-specific CD8^+^ T cells among Vβ12^-^ cells in tumors of NP-PEG-BayK8644-treated mice comparing progressor and responder animals. * p < 0.05, Student’s *t* test. **k)** Absolute numbers of OVA-specific CD8^+^ T cells among Vβ12^-^ cells in tumors of NP-PEG-BayK8644-treated mice comparing progressor and responder animals. * p < 0.05, Student’s *t* test. **l)** Representative flow cytometry dot plots of OVA-specific CD8^+^ T cells in tumors of NP-PEG-BayK8644-treated mice comparing progressor and responder animals.

## Conclusions

High TMEM176A/B expression has been associated with diminished overall survival in different human tumors, suggesting that it is a potential pharmacological target to control cancer progression [17,20,21]. However, controversial data have been reported concerning the role of this cation channel in controlling anti-tumoral CD8^+^ T cell responses. Jiang et al. reported that in B16 melanoma, tumor growth was accelerated in *Tmem176b*^*-/-*^ mice in association with diminished tumor infiltration by CD8^+^ T cells in comparison to WT animals [31]. In contrast, we have shown that *Tmem176b*^*-/-*^ mice control EG7 lymphoma, LL2 lung and MC38 colon cancer growth in an inflammasome and CD8^+^ T cell-dependent manner. Moreover, pharmacological blockade of TMEM176B with BayK8644 triggers *Tmem176b*, inflammasome and CD8^+^ T cell-dependent anti-tumoral immune responses leading to tumor control in preventive but not in therapeutic protocols [17]. Thus, acknowledging the dual role played by TMEM176B in controlling inflammasome activation and antigen cross-presentation and its impacts in different models is key to better understand this controversy as well as to improve the therapeutic potential of BayK8644.

Here we report that formulating the TMEM176A/B inhibitor BayK8644 in chitosan nanoparticles dissociated the dual role of TMEM176B on innate and adaptive immunity. Thus, NP-PEG-BayK8644 prevented inhibition of cross-presentation by the free compound while triggering inflammasome activation. Furthermore, NP-PEG-BayK8644 improved the inflammasome-activating capacity of BayK8644 in *in vitro* studies.

NPs are promising strategies for the controlled release of drugs, particularly in oncology [32]. Although chitosan is considered a safe polymer, variations in origin, composition and molecular weight among other parameters have complicated the translational pathway of chitosan nanoparticles [33]. Further research will need to determine whether encapsulation of the compound in NPs of other materials may give similar results in preclinical models.

In conclusion, here we add preclinical proof-of-concept of a nanotechnology-based method that improves the anti-tumoral therapeutic efficacy of BayK8644 by uncoupling the dual role of TMEM176B on innate and adaptive immunity.

## Acknowledgments

This work was supported by Uruguay INNOVA-2 from ANII, PEDECIBA, ECOS-SUD and FOCEM (MERCOSUR Structural Convergence Fund) COF 03/11 grants to MH.

---

**Supplementary figure 1.**
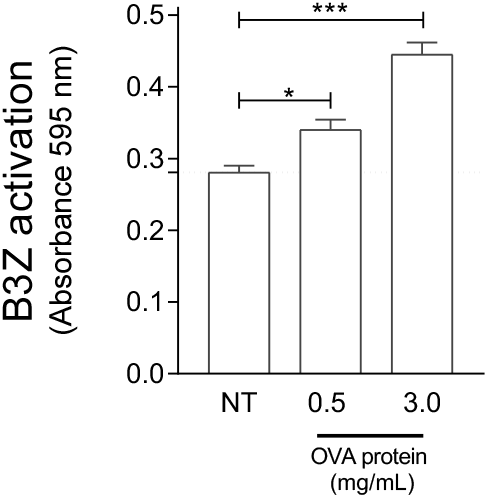
Cross-presentation assay. BMDCs were loaded for 45 minutes with different doses of OVA protein or left untreated (NT). After washing, cells were co-cultured with B3Z CD8^+^ T cells. T cell activation was studied through colorimetry as detailed in supplementary Methods. One experiment representative of four is shown. One-way ANOVA test. * p< 0.05; *** p<0.001.

**Supplementary figure 2.**
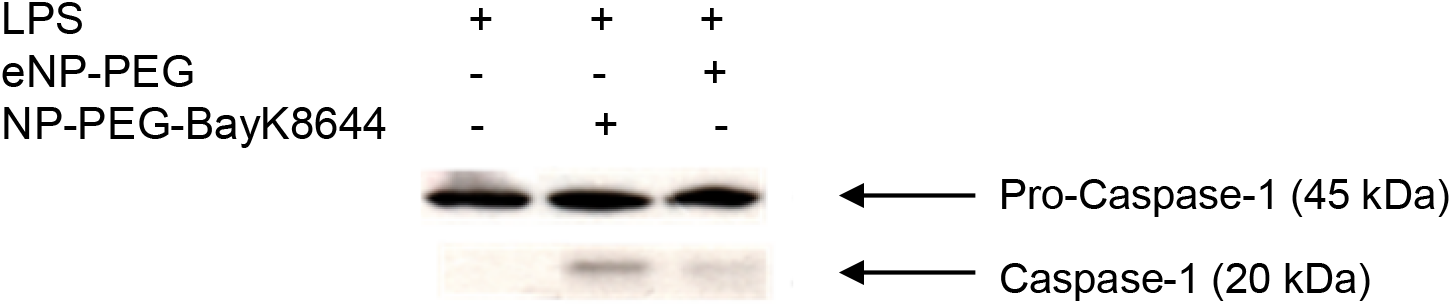
Western blot analysis of Caspase-1 activation. BMDCs were treated for 4 hr with LPS and then incubated for 16 hr with eNP-PEG or NP-PEG-BayK8644. Pro-Caspase-1 and Caspase-1 were assessed in the cell lystae and culture supernatant respectively. One experiment representative of two is shown.

**Supplementary figure 3.**
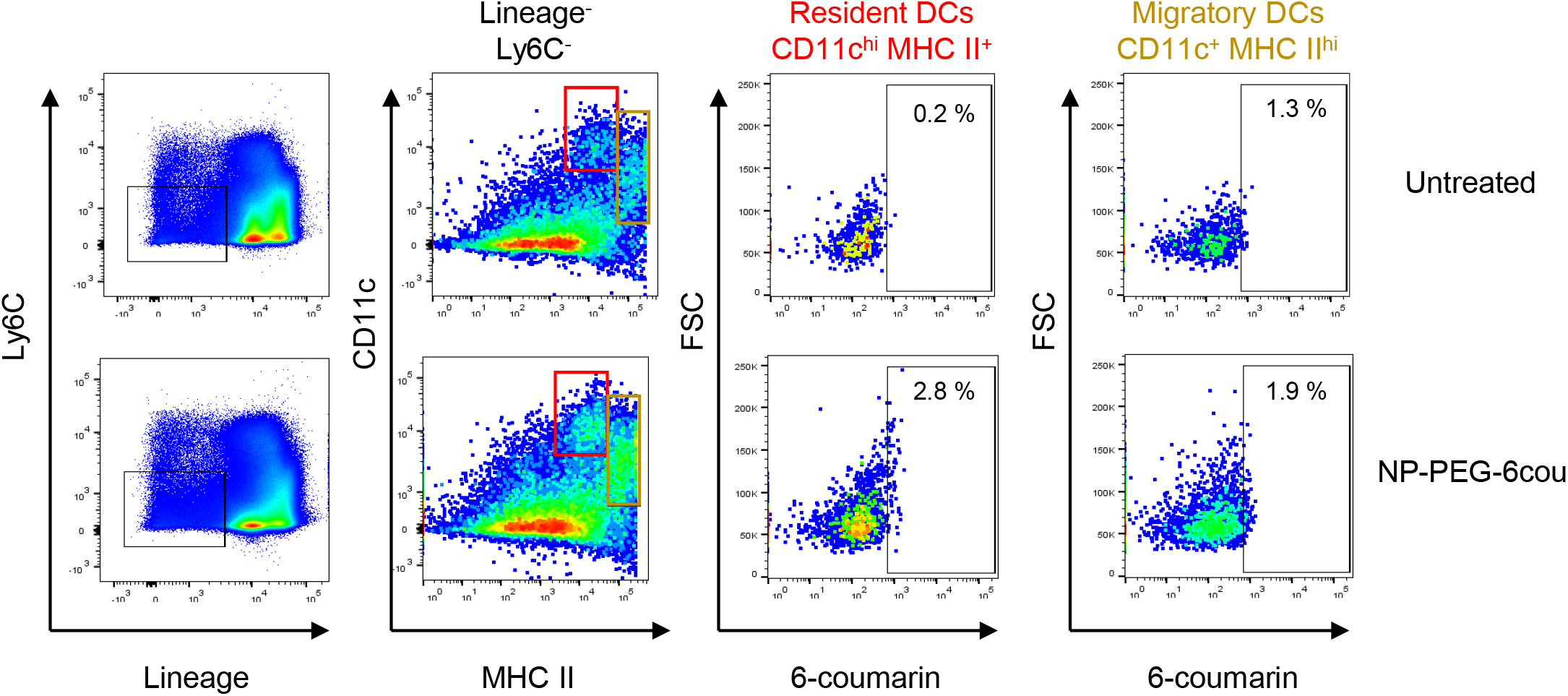
In vivo tracking of intratumoral injected fluorescent nanoparticles. WT mice bearing EG7 tumors were injected i.t. with fluorescent NPs (NP-PEG-6-cou). Twenty-four hr later, the tumor-draining lymph nodes were harvested. Cell suspensions were stained with the indicated antibodies and analyzed by flow cytometry. Lineage: anti-B220, anti-TCRb, anti-NK1.1. Viable cells were determined by forward scatter-side scatter dot plot and by doublet exclusion the DC population was analyzed considering T-cell, B-cell and NK-cell exclusion. Within the MHC-II^+^ and CD11c^+^ population, the percentage of migratory and residents DCs was determined and then the percentage of 6-coumarin^+^ cells was analyzed within those populations. One animal representative of two untreated and two injected with NP-PEG-6cou are shown.

**Supplementary table 1.**
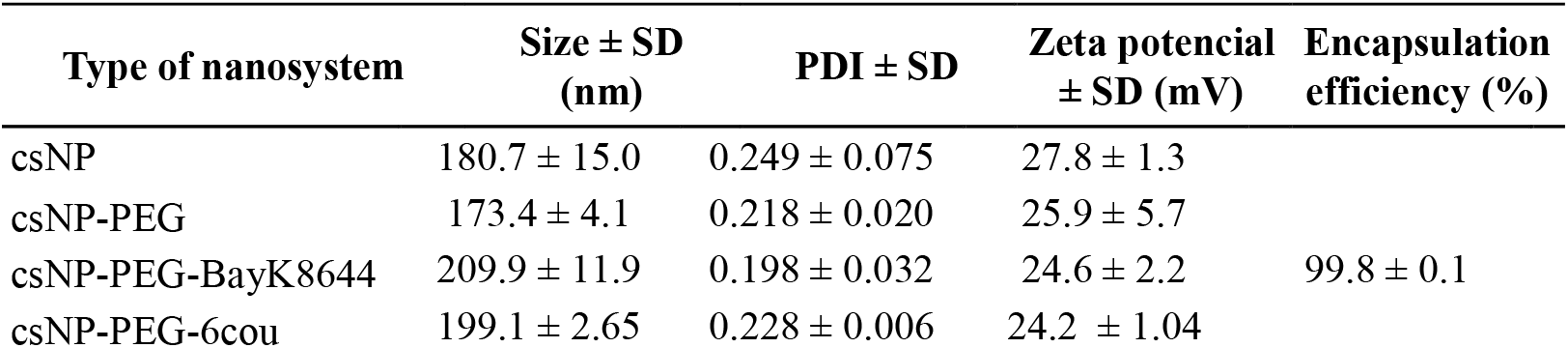
Physico-chemical properties of the nanosystems obtained (mean ± s.d.; n=3)

